# Neuroepithelial reprogramming and ERBB vulnerability in canine acanthomatous ameloblastoma

**DOI:** 10.64898/2026.03.03.709191

**Authors:** Andreas Stephanou, Bo Shui, Deanna Mische, Michael Byron, William P. Katt, Marina Chan, Jennifer K. Grenier, Iwijn De Vlaminck, Gerald E. Duhamel, Taranjit S. Gujral, Praveen Sethupathy, Santiago Peralta

**Affiliations:** Department of Biomedical and Translational Sciences, College of Veterinary Medicine, Cornell University, Ithaca, NY 14853, USA; Nancy E. and Peter C. Meinig School of Biomedical Engineering, Cornell University, Ithaca, NY 14853, USA.; Human Biology Division, Fred Hutchinson Cancer Center, Seattle, WA 98109; Department of Clinical Sciences, Clinical Programs Center, College of Veterinary Medicine, Cornell University, Box 31, Ithaca, NY 14853, USA; Biotechnology Resource Center, Cornell University, Ithaca, NY 14853, USA

**Author notes:** **Correspondence should be addressed to:** P.S. and S.P. These authors contributed equally. **Author Contributions:**.

**Keywords:** Single-nucleus RNA sequencing, Ameloblastoma, ERBB signaling

## Abstract

Canine acanthomatous ameloblastoma (CAA) is a locally invasive benign oral neoplasm that is difficult to distinguish from canine oral squamous cell carcinoma (COSCC) due to overlapping clinical, radiologic, and histologic features. Although both tumors exhibit MAPK pathway activation, their mutational landscapes are distinct. Furthermore, previous studies using bulk RNA sequencing (RNA-seq) have demonstrated pronounced differences in programs related to, among others, hypoxia, PI3K-AKT signaling, and cell proliferation. However, these bulk studies lacked the resolution to elucidate the cellular heterogeneity of CAA relative to COSCC. We therefore performed single-nucleus RNA-seq to define the cellular gene expression landscape of CAA, COSCC, and healthy gingiva. Across ∼205,000 nuclei, we identified major epithelial, immune, endothelial, and mesenchymal populations, as well as two epithelial subtypes uniquely enriched in CAA. The CAA-specific keratinocytes exhibited a neuronal-like expression program defined by synaptic regulators, KRAS-associated signaling pathways, and markedly elevated expression of *PEG3, ERBB4, GABRB1, MAGI2*, and *CASK*. These findings were validated by bulk RNA-seq, qPCR, and immunohistochemistry, which demonstrated strong nuclear localization of PEG3 exclusively in CAA epithelium. A kinase inhibitor screen independently identified ERBB4 as a candidate therapeutic vulnerability, and pharmacologic inhibition with neratinib was effective. Together, these findings reveal a previously unrecognized neuroepithelial cell state that defines CAA, distinguishes it from COSCC, and reveals unique diagnostic and therapeutic signaling dependencies. Given the molecular and histopathologic parallels between CAA and human ameloblastoma, these data further position CAA as a naturally occurring comparative model for studying ameloblastoma biology and therapeutic vulnerabilities.

**Significance Statement:** Canine acanthomatous ameloblastoma (CAA) is a common locally aggressive oral neoplasm often misdiagnosed as canine oral squamous cell carcinoma (COSCC) due to overlapping clinical, radiologic, and histologic features. Using single-nucleus RNA sequencing of CAA, COSCC, and healthy gingiva, we resolve the microenvironment of these tumors and detect two epithelial subpopulations unique to CAA. These CAA-specific keratinocytes exhibit a neuroepithelial-like signature characterized by synaptic regulators, KRAS-associated signaling, and elevated PEG3 and ERBB4 expression. Functional kinase inhibitor screening of patient derived microtumor slices independently converged on ERBB4 as an effective therapeutic target. Together, our study provides a molecular framework for distinguishing CAA from COSCC and establishes a comparative oncology model with translational relevance to human ameloblastoma and other ERBB-driven epithelial neoplasms.

## Introduction

Canine acanthomatous ameloblastoma (CAA) is a common benign odontogenic epithelial neoplasm of the oral cavity^1,2^. It has been reported in various dog breeds of a wide age range (mean 9 years old) with no apparent sex bias^1,2^. Despite its lack of metastatic potential, it is locally invasive, often infiltrating the surrounding tissue, including the jawbone^3,4^. The most effective current treatment is wide-margin excision (i.e., mandibulectomy or maxillectomy) that is associated with low recurrence rates, but significant functional consequences^5–8^. Other more conservative interventions, such as radiation therapy, marginal excision, and intralesional bleomycin injection, may lead to greater function and cosmetic outcome, but have substantially higher recurrence and lower long-term remission rates^9–11^. Although less prevalent compared to dogs, ameloblastoma also occurs in people. Moreover, the biologic and clinical behavior exhibited by CAA and human ameloblastoma are comparable including low proliferation activity and similar transcriptional signatures, suggesting that CAA may be an excellent and important model system for cross-species studies that attempt to develop better treatment approaches^4,12–17^.

A significant difficulty in diagnosing and treating CAA has been to differentiate it from a second common oral malady in dogs, canine oral squamous cell carcinoma (COSCC). Histopathological examination of biopsy specimens is the gold standard approach for definitive identification of either disease^18^. However, CAA shares many clinical, radiological, and histological features with COSCC, which often leads to misdiagnosis (∼30% of cases)^2,19^. Accurate distinction is critical given the clinical implications, as COSCC exhibits metastatic potential and markedly faster growth^4,20^.

Molecular characterization of CAA and COSCC faces similar challenges. Both CAA and COSCC have an overactive MAPK pathway, but their mutational landscape is distinct^16,17,21^. Indeed, over 95% of CAA tumors harbor an *HRAS* p.Q61R somatic mutation, whereas COSCC tumors harbor other MAPK pathway activating mutations (i.e., *BRAF* p.V595E, *HRAS* p.Q61L) or heretofore uncharacterized mutations that may or not dysregulate the MAPK pathway^21,22^. Transcriptomic analyses have further revealed pronounced differences: COSCC is enriched for gene programs related to hypoxia, PI3K-AKT signaling, and cell proliferation, while expression of *ODAM* (a gene involved in tooth development and enamel maturation) has emerged as a potential CAA-specific marker^16^.

The molecular characterization of CAA and COSCC to date has been based on bulk sequencing technologies^16,17^. As a result, the cellular composition and molecular heterogeneity of CAA tumors remain unexplored. Moreover, any subtle signaling emerging from rare, specialized cell types would likely be drowned out in the bulk signal, rendering it undetectable. These limitations have stymied progress on defining detailed molecular and cellular features of CAA that distinguish it from COSCC. Fortunately, recent advances in genome technologies facilitate large-scale single-cell/single-nucleus gene expression profiling of hundreds of thousands of individual cells, making possible the dissection of the heterogeneous microenvironment that drives various diseases^23,24^.

The goal of this study was to characterize the gene expression landscapes of CAA and COSCC at unprecedented resolution, using single-nucleus technology, to be able to define the unique cellular composition and molecular features of either disease. To provide comparative biological context, we used healthy gingival tissue (HGIN) as a normal control. Here, we describe our findings showing that CAA tumors have a very large population of neuronal-like cells, marked by *PEG3*, and that CAA tissue can be effectively targeted using neratinib, a potent ERBB and MEK inhibitor^25^. These findings may have significant ramifications for both accurately diagnosing CAA and potentially developing new treatment strategies, as well as further informing comparative features of relevance in human oncology.

## Results

### SnRNA-seq reveals unique CAA-specific cell types in CAA

To investigate the cellular heterogeneity of CAA, we performed single-nucleus RNA sequencing (snRNA-seq) on 19 samples: six CAA, ten COSCC, and three adjacent HGIN (**Fig. 1A**). After quality control, including low cell count filtering and doublet removal (**Methods**), we were left with 204,967 high-quality nuclei (**Fig. 1A**). The resulting nuclei had a median of 5,778 total counts (HGIN:2,240; COSCC: 7,250; and CAA: 4,843), a median of 2,496 non-zero expressing genes (HGIN:1,355; COSCC: 2,791; and CAA: 2,289), and a median of 0% mitochondrial counts across all conditions (**Fig. 1B**). To determine whether any samples were mislabeled, we generated pseudobulk samples from the snRNA-seq data by summing gene counts for each patient sample. We then performed principal component analysis (PCA) to cluster the pseudobulk data. The pseudobulk samples separated well by condition, confirming that no samples were mislabeled and that the between-condition variance was much greater than the within-condition variance along the top principal components, consistent with expectation (**Fig. 1C**). To gain further confidence in the reliability of the snRNA-seq data, we asked whether the gene expression relationship between CAA and COSCC was consistent with a newly generated bulk RNA-seq dataset (DS1) and a previously published one (DS2^16^) (**Supp. Fig. 1A-C**). To do so, we performed differential gene expression analysis (DGEA) on the pseudobulk data against the two bulk RNA-seq datasets. The gene expression differences between the datasets were strongly correlated with a Pearson r = 0.73 (**Fig. 1D**). Hence, overall, we concluded that the snRNA-seq data are reliable.

**Figure 1:**
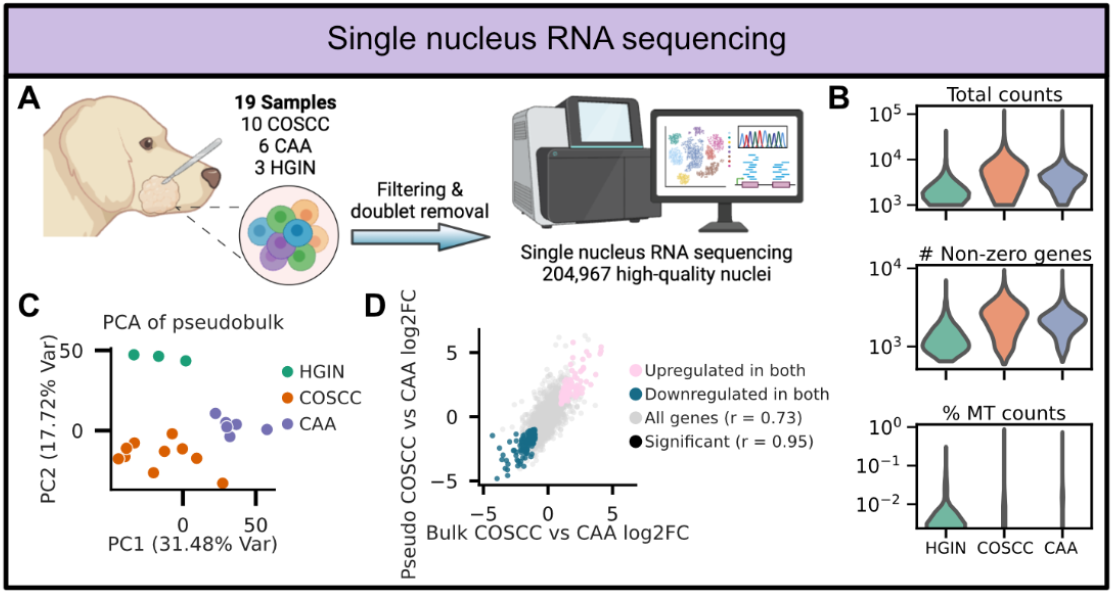
Workflow and quality assessment of snRNA-seq data. **A)** Experimental workflow of CAA, COSCC, and HGIN snRNA-seq. Created with BioRender.com. **B)** Violin plots showing the i) total counts (reads), ii) number of non-zero expression genes, and iii) percentage of mitochondrial counts per cell. **C)** PCA plot of generated pseudobulk data using the top 2000 most highly variable genes. **D)** Scatterplot of overlapped bulk (x axis) and pseudobulk (y axis) DGE analysis between CAA and COSCC conditions. Colored spots represent significantly (|log2FC|>1 and adjusted p < 0.01) upregulated (pink) and downregulated (teal) genes for both dataset DGEs.

We next clustered and annotated the nuclei by major cell type categories. These included epithelial, immune, endothelial, and mesenchymal cells, which were classified using canonical markers (**Fig 2A**). As expected, epithelial cells dominated across all conditions, comprising over 70% of all cells in the tumoral conditions and over 90% of the healthy gingiva (**Fig 2A**). Notably, we observed an expansion of non-epithelial cell types, such as endothelial cells and immune cells, in the tumor microenvironments (TMEs) relative to healthy gingiva. Next, we determined the cellular subtypes within each major cell category, namely mesenchymal, immune, endothelial, and epithelial cells. Considering mesenchymal cells, we detected melanoblasts only in the healthy gingiva. However, a substantial population of fibroblasts as well as vascular smooth muscle cells and pericytes were found in the TMEs for both CAA and COSCC samples (**Fig. 2B**). The immune repertoire was markedly altered in the TME, with an increased proportion of most immune subtypes, such as M2 macrophages, CD4+ and CD8+ T cells, compared to HGIN (**Fig. 2C**). Additionally, we observed evidence of vascularization in the TMEs, indicated by an increased presence of capillary and venous endothelial cells (**Fig. 2D**). Upon examining the epithelial cell subtypes, which are presumed to give rise to the tumor cells in CAA and COSCC, we detected two epithelial subtypes that were exclusive to the CAA tumors: (i) CAA-specific junctional keratinocytes (KCs) and (ii) CAA-specific basal KCs (**Fig. 2E**). Furthermore, COSCC contained a substantial population of proliferating basal KCs, consistent with previous reports of an elevated Ki-67 index (**Fig. 2E**). Collectively, our results indicate that CAA is characterized by unique epithelial subtypes and profound remodeling of the tumor microenvironment.

**Figure 2:**
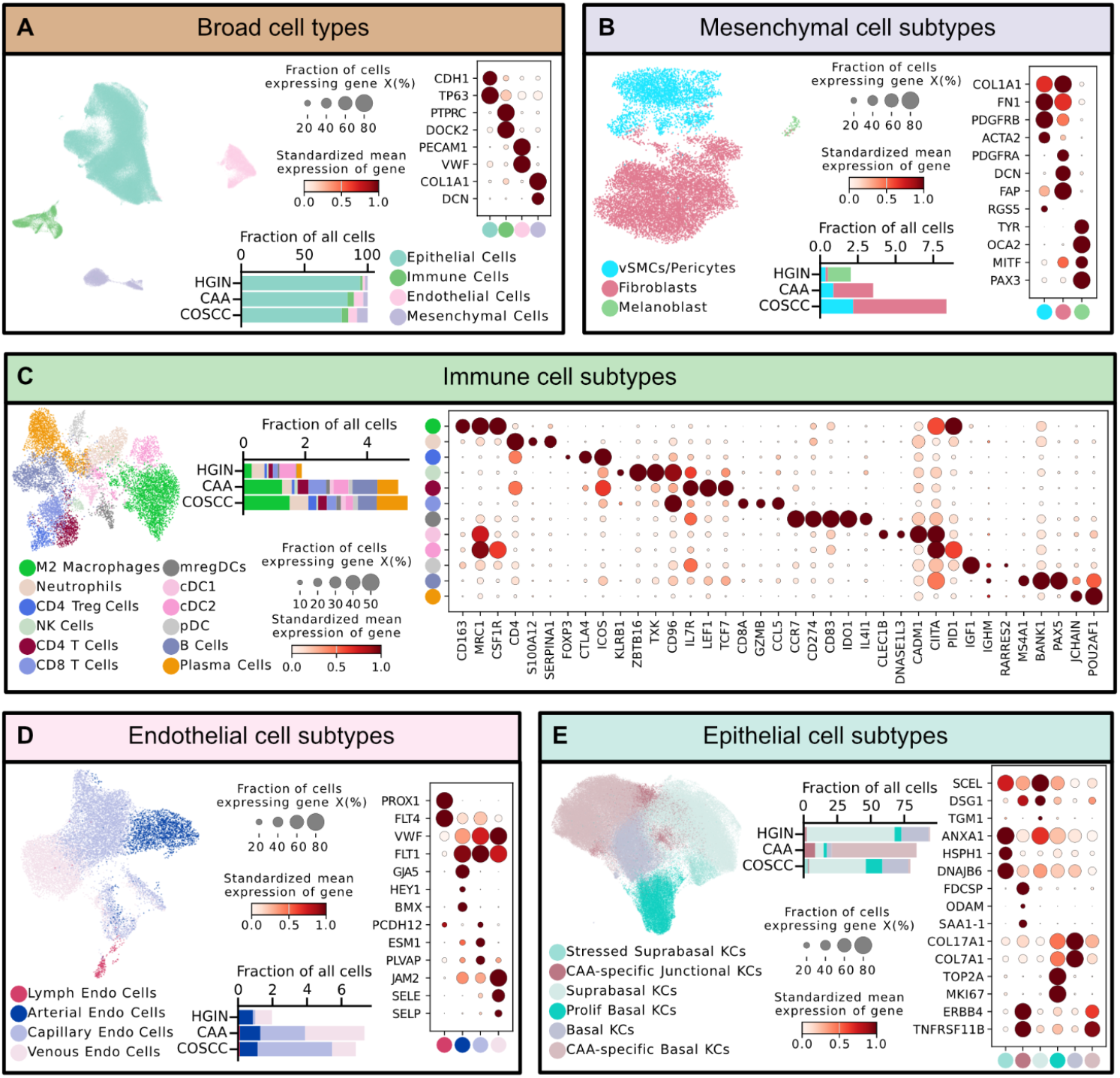
Characterization of cellular subtype diversity captured by snRNA-seq. **A)** The complete dataset was first annotated into broad cell type classes. These annotations were then further resolved to identify finer subpopulations within each major compartment, including **B)** mesenchymal, **C)** immune, **D)** endothelial, and **E)** epithelial cell subtypes. Uniform Manifold Approximation and Projection (UMAP) plots colored by cell type and key marker genes are shown for each analysis. In the dot plots, dot size represents the percentage of cells expressing each gene (non-zero expression), and dot color reflects the standardized mean expression. Bar plots display the fraction of total cells contributed by each subtype per condition.

### CAA-specific keratinocytes (CAA KCs) exhibit a neuronal-like signature

To better characterize the CAA KCs, we performed pairwise DGEA between each CAA KC subtype and all other epithelial cell types as well as the merged TME (**Methods**). This resulted in 168 significantly upregulated genes (adjusted p < 0.01 and log_2_FC > 2) in either of the CAA KCs, with the top 20 most highly expressed of these shown in **Fig. 3A**. These 168 genes were defined as the CAA-specific KC signature, which when used in a PCA, effectively separated CAA from COSCC and HGIN, not only using the pseudobulk data but also the CAA bulk RNA-seq datasets (DS1 and DS2) (**Fig. 3B**). Next, we sought to describe the CAA-specific KC signature using gene expression overrepresentation analysis. Using the MSigDB^26–28^ C5 (GO gene sets) and C6 (oncogenic signatures) collections, the top significantly enriched terms were associated with synaptic and neuronal function, as well as the KRAS signaling pathway (**Fig. 3C**). The genes contributing to these enriched terms are shown in **Fig. 3D**.

**Figure 3:**
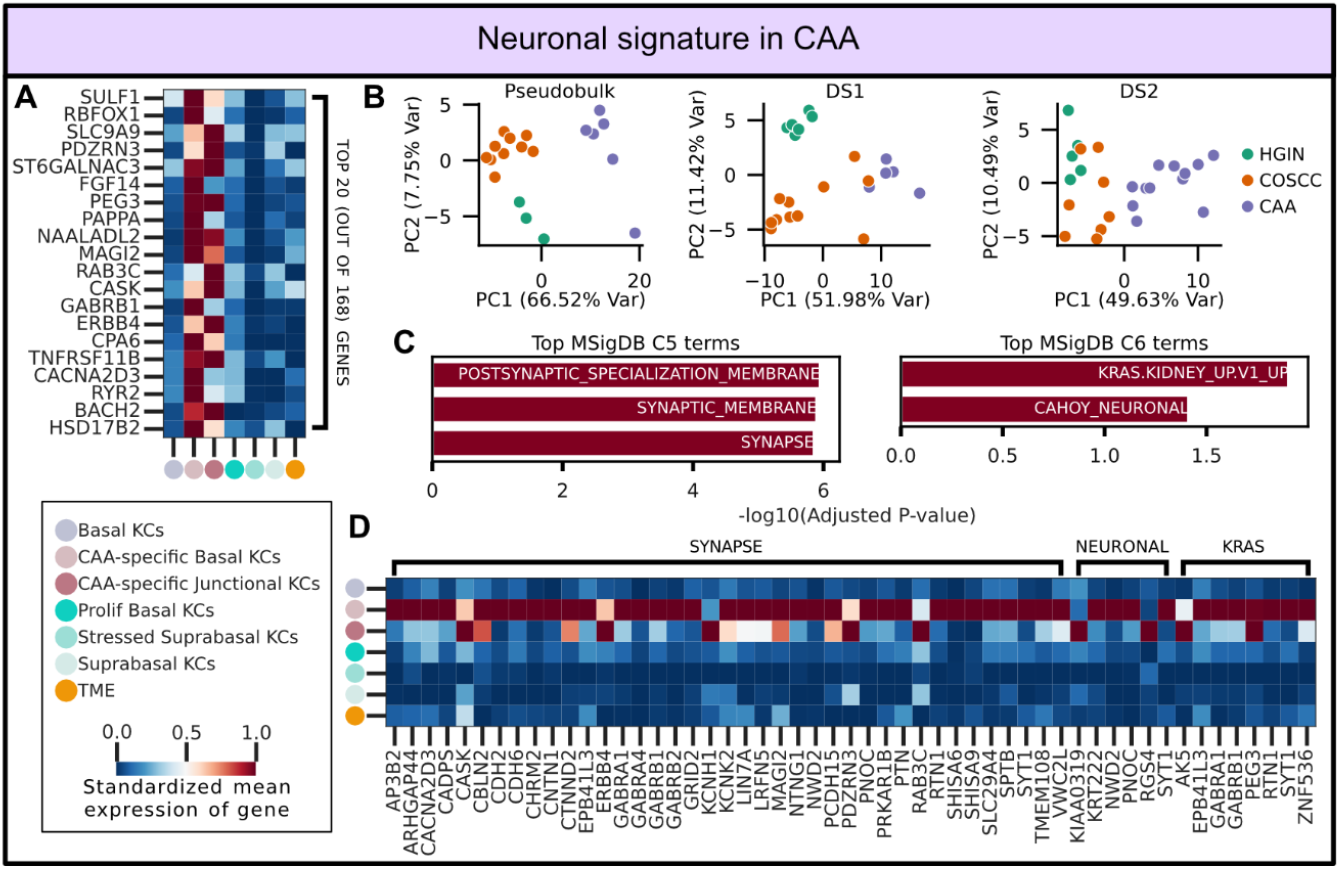
Characterization of neuronal signature detected in CAA-specific KCs. **A)** Heatmap of the top 20 (out of 168) most highly expressed DEGs in the CAA-specific KCs. **B)** PCA plots of pseudobulk and bulk (DS1 and DS2) RNA-seq samples using the CAA-specific KC signature. **C)** Top significant terms from gene expression overrepresentation analysis of CAA-specific KC signature using MSigDB’s C5 (GO gene sets) and C6 (oncogenic signatures) collections. **D)** Heatmap of the genes contributing to the MSigDB terms SYNAPSE, CAHOY_NEURONAL, and KRAS.KIDNEY_UP.V1_UP.

To validate the CAA-specific KC signature, we selected candidate genes and evaluated their expression across datasets and modalities. Candidate selection was based on the overlap of (1) the top 20 genes of the CAA-specific KC signature, (2) genes associated with synaptic, neuronal, and KRAS terms, and (3) significantly upregulated genes from the pseudobulk CAA vs COSCC DGEA (**Fig. 4A**). Ranking these overlapping genes by log_2_(fold-change) yielded five top candidates: *PEG3, GABRB1, ERBB4, MAGI2*, and *CASK* (**Fig. 4A**). We focused our validation on the top candidate, *PEG3*, and on *ERBB4*, as previous work has described its critical role in the human analog of CAA^29^. In both bulk RNA-seq datasets, *PEG3* was significantly upregulated in CAA compared to both HGIN and COSCC (Kruskal–Wallis test with post-hoc Dunn’s test and FDR correction; adjusted p < 0.05) (**Fig. 4B**). *ERBB4* was not detected in dataset 2 (DS2) but was significantly upregulated in CAA relative to HGIN and COSCC in dataset 1 (DS1) (Kruskal–Wallis test with post-hoc Dunn’s test and FDR correction; adjusted p < 0.05) (**Fig. 4B**). Orthogonal validation of *PEG3* and *ERBB4* using qPCR further confirmed that *PEG3* and *ERBB4* are transcribed at markedly higher levels in CAA as compared to COSCC (**Fig. 4C**).

**Figure 4:**
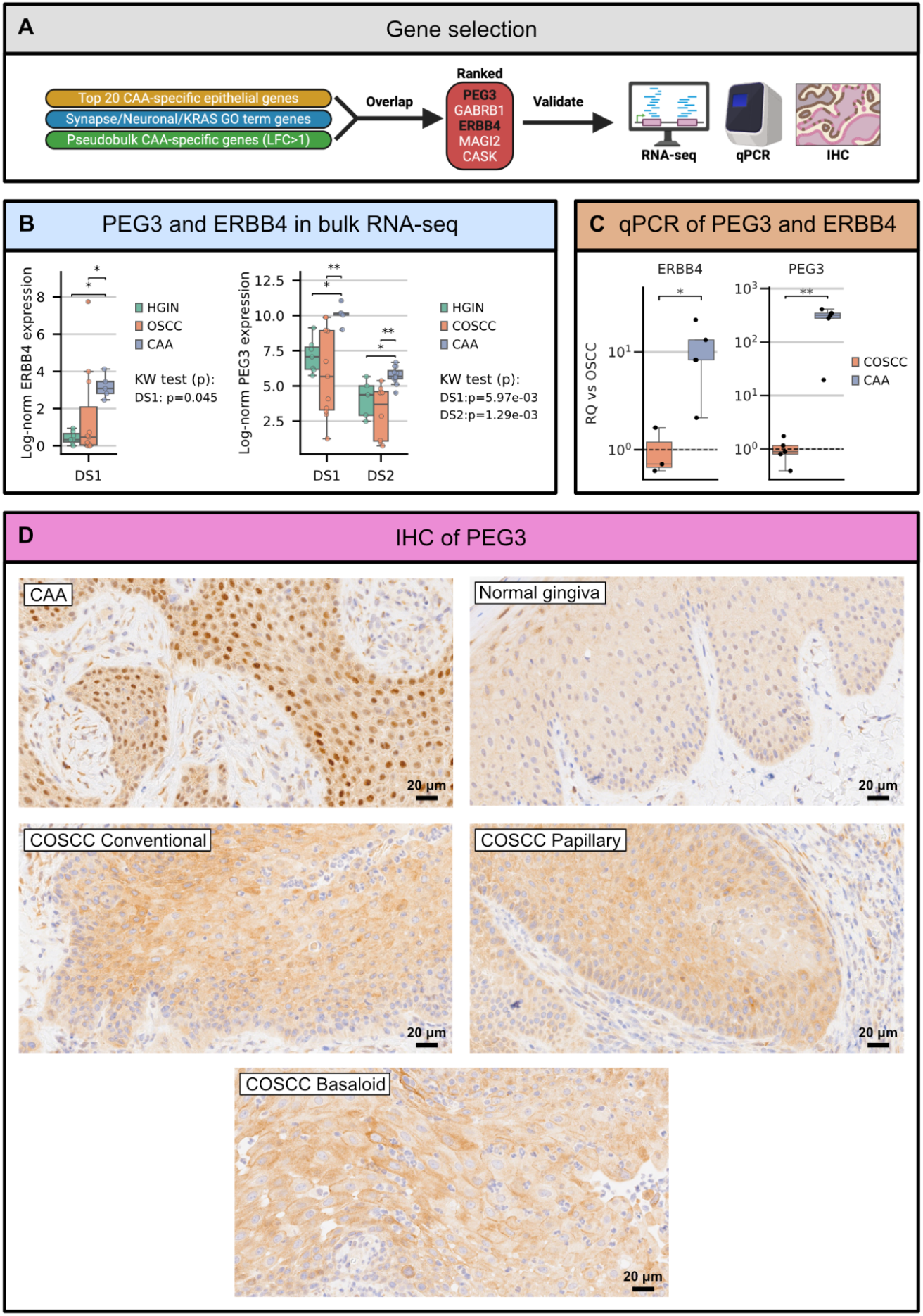
Validation of *PEG3* and *ERBB4* expression in CAA using various modalities. **A)** Workflow schematic of candidate gene selection. Created with BioRender.com. **B)** Boxplots of *PEG3* and *ERBB4* log normalized expression across samples in bulk RNA-seq datasets DS1 and DS2. Boxes are colored based on tumor status and pairwise comparison adjusted p-values are indicated by * < 0.05 and ** < 0.01. **C)** Box plots colored by tumor status showing qPCR relative quantification (RQ) values for *ERBB4* and *PEG3*. Pairwise comparison adjusted p-values are indicated by * < 0.05 and ** < 0.01. **D)** Immunohistochemistry of representative formalin-fixed and paraffin-embedded tissue sections of CAA, healthy gingiva, COSCC conventional, papillary, and basaloid subtypes with PEG3-specific polyclonal rabbit IgG antibody. Scale bar, 20 µm.

Immunohistochemistry assessment of PEG3 protein expression revealed variable expression between CAA and COSCC. While control gingiva showed non-specific immunoreactivity (**Fig. 4D**), PEG3 was differentially expressed in CAAs compared with COSCC subtypes. In CAAs PEG3 was strongly localized to the nucleus of all neoplastic epithelial cells with fine stippling along the cytoplasmic membrane (**Fig. 4D**). By contrast, in the three COSCC subtypes, PEG3 immunoreactivity was weaker and limited to coarse cytoplasmic aggregates that extended to the cytoplasmic membrane (**Fig. 4D**).

### Polypharmacology-based kinome screening identifies ERBB4-centered signaling as a shared functional dependency in CAA

Although our snRNA-seq analysis revealed a distinct CAA-specific epithelial population enriched for ERBB4 and MAPK-associated signaling programs, transcriptional enrichment alone does not establish functional dependency. Given the extensive redundancy and compensatory signaling within kinase networks, we implemented an unbiased functional approach to determine which kinases are required for CAA tumor viability. To address this, we performed a polypharmacology-based kinome screen using Kinase inhibitor Regularization (KiR) (**Fig. 5A**)^30,31^. KiR leverages the fact that most kinase inhibitors are not highly selective but instead target multiple kinases. Rather than viewing this as a limitation, KiR exploits polyselectivity to deconvolve kinase dependencies computationally. Our curated panel of 32 kinase inhibitors collectively covers over 90% of the kinome, enabling broad functional interrogation of signaling networks. By integrating drug response profiles with known kinase-inhibitor interaction matrices through machine learning, KiR identifies kinases most likely driving a phenotype while accounting for redundancy and compensatory pathways often missed by single-agent approaches^30,31^. This method has been extensively validated across diverse biological systems, including cancer growth and migration, cytokine signaling, malaria, and viral infection models^32–38^.

**Figure 5:**
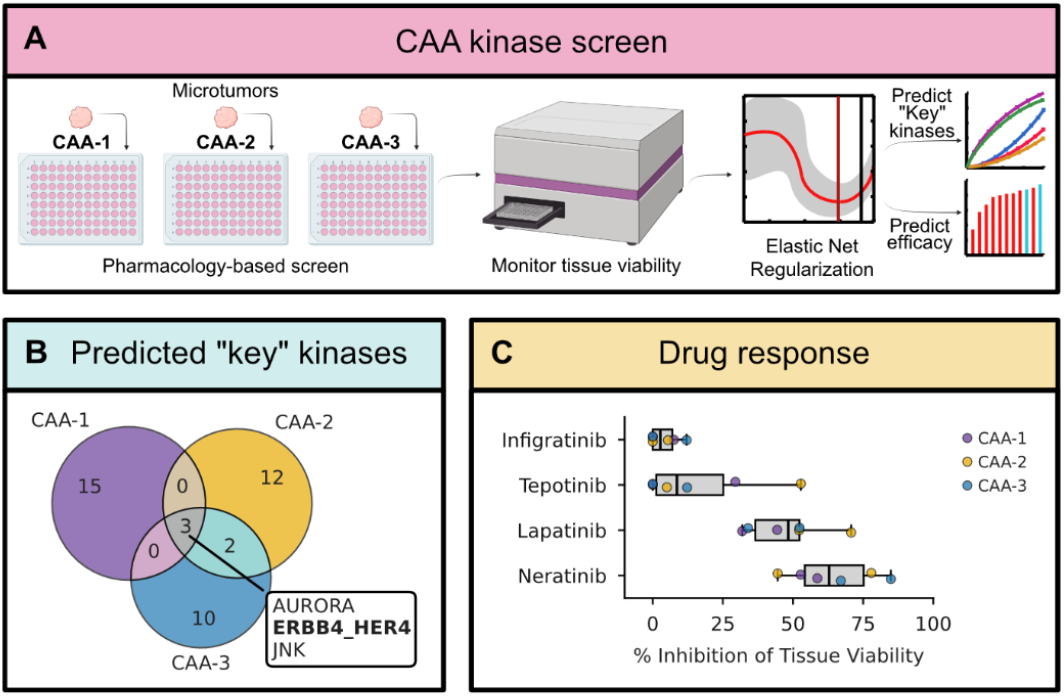
Functional kinase screening and drug response profiling in CAA tumor slices. **A)** Kinase screen method overview on 3 CAA patients. Created with BioRender.com. **B)** Venn-diagram of predicted “key” kinases for each CAA patient. Kinases predicted in all 3 CAA patients are shown. **C)** Box plots showing the % inhibition of cell viability by the FDA approved therapeutic drugs infigratinib (FGFR inhibitor), tepotinib (MET inhibitor), lapatinib (high affinity ERBB1/2, lower affinity ERBB4 inhibitor), and neratinib (pan-ERBB higher affinity ERBB4 inhibitor, and MEK inhibitor).

We applied this approach to three independent CAA cases using patient-derived microtumor models^39^. Microtumors were treated with DMSO vehicle or each of the 32 kinase inhibitors, and tissue viability was monitored over five days. The resulting viability profiles were used to construct case-specific KiR models, which ranked predicted kinase dependencies for each tumor. Across all three CAA cases, KiR identified both shared and case-specific kinase vulnerabilities (**Fig. 5B**). Notably, three kinases emerged as common predicted dependencies in all three cases: ERBB4, Aurora, and JNK. In addition, partial overlaps were observed, including predicted vulnerability to ARAF and ACK1 in both CAA-2 and CAA-3, suggesting possible convergence on MAPK-associated circuitry in a subset of CAA tumors.

Importantly, ERBB4 was independently identified as a top dependency across all three cases, converging with our single-nucleus transcriptomic data that showed *ERBB4* enrichment within CAA-specific epithelial populations. Functional validation further supported this prediction: treatment with neratinib (1 µM), an irreversible ERBB-family and MEK inhibitor, produced strong inhibition of viability in all three CAA microtumor models (**Fig. 5C**). The magnitude of response exceeded that observed with lapatinib, a high affinity ERBB1/2 but lower affinity ERBB4 inhibitor, suggesting that ERBB4 targeting is particularly critical in treating CAA (**Fig. 5C**). As FGFR and MET signalling are not predicted to be implicated in CAA progression, tepotinib and infigratinib were used as negative controls and indeed showed minimal response (**Fig. 5C**).

Together, these findings demonstrate that integrating polypharmacology-based kinome screening with computational network modeling identifies recurrent kinase dependencies in CAA and functionally validates ERBB4-centered signaling as a shared and actionable vulnerability.

## Discussion

In this study, we provide the first snRNA-seq atlas of CAA and demonstrate that it is defined by a unique neuroepithelial cell type signature absent in COSCC and healthy gingiva. Across ∼205,000 nuclei we detected two epithelial subpopulations that were unique to CAA and which expressed a neuronal-like signature. This signature was composed of genes associated with synaptic and neuronal function, as well as the KRAS signaling pathway. Of these genes, *PEG3, GABRB1, ERBB4, MAGI2*, and *CASK* showed a markedly elevated expression, and PEG3 protein localized predominantly to the nucleus, consistent with its role as an activated transcription factor. In contrast, PEG3 was largely absent from the nucleus in COSCC and healthy gingiva. Most importantly, by orthogonal approach, our CAA kinase screen revealed ERBB4 as a potential therapeutic target which was validated with the use of neratinib. These findings expand substantially on prior bulk RNA-seq studies by revealing the molecular heterogeneity of CAA at single-cell resolution and detecting a previously unidentified neuronal-like signature marking the CAA neoplasm which can be targeted therapeutically by a potent ERBB-family and MEK inhibitor. By delineating these molecularly distinct epithelial populations, our work refines the biological understanding of CAA and highlights features that have important implications for diagnosis, comparative oncology, and therapeutic targeting.

The work we present is consistent with, and significantly expands upon, previous studies in both canine and human ameloblastoma. Prior bulk RNA-seq studies demonstrated that CAA and human ameloblastoma share a low proliferative ability and the expression of the odontogenic marker *ODAM*, which indicated conservation across species^16^. However, these bulk approaches were limited in resolution and did not capture the cellular heterogeneity of these conditions. Our single-nucleus analysis overcomes this limitation and reveals that the observed *ODAM* enrichment in CAA is limited to a discrete subset of the CAA epithelium, which is absent in COSCC and healthy gingiva. Additionally, the expression of *ERBB4* in CAA-specific epithelial cells reinforces findings in human ameloblastoma in which ERBB family signaling was supportive of tumor progression and reinforces the translational potential of our work^29^. The presence of synaptic regulators such as *PEG3, MAGI2*, and *CASK* further suggests that CAA tumor cells may be capable of bidirectional communication with peripheral neurons, consistent with prior work showing that human ameloblastomas are innervated and that afferent nerve fibers enter in close contact with the tumor epithelium^40^. Thus, our study bridges key findings in canine and human ameloblastoma and extends prior transcriptomic work.

The discovery of neuronal-like KCs in CAA has significant therapeutic potential. The enriched expression of KRAS genes, *ERBB4*, and synaptic regulators suggests that CAA may engage in atypical MAPK signaling distinct from COSCC, which could reveal novel therapeutic vulnerabilities within ERBB or neurotrophic signaling axes. Furthermore, since CAA is characterized as an HRAS-driven tumor, it may serve as a valuable comparative model for investigating RAS-dependent epithelial plasticity and tumor–nerve interactions across cancers.

While single nucleus technologies allow for high-resolution profiling of frozen tissue samples, they do bias the captured transcriptome toward nuclear RNA and are insensitive to cytoplasmic transcripts, including key neuronal and synaptic genes. Furthermore, our access to healthy tissue was limited, reducing our ability to fully resolve the normal tissue cellular diversity. Moreover, following cellular dissociation, spatial information is lost which limits our ability to characterize the neuronal-like epithelial cells in their native context. Future studies could employ spatial transcriptomics to address this limitation. Lastly, additional functional work, such as KRAS inhibition, could strengthen our mechanistic conclusions. Together, our work expands the molecular framework of CAA and uncovers a previously unrecognized neuroepithelial program and likely therapeutic vulnerability that reshapes how this tumor is classified, studied, and targeted.

## Materials and Methods

### Sample collection

Tissue samples corresponded to cryopreserved material collected from client-owned dogs undergoing standard-of-care diagnostic or therapeutic interventions at the Cornell University Hospital for Animals. Sample collection was done in accordance with protocols previously reviewed and approved by Cornell University’s Institutional Animal Care and Use Committee (protocols #2015-0117 and #2022-0166). Signed informed consent was obtained from all dog owners prior to experimental sample collection. Tumor or normal tissue samples were obtained using a 2- or a 4-mm punch while animals were under general anesthesia following standard clinical protocols. Sample collection and general anesthesia were performed or directly supervised by board-certified specialists in veterinary dentistry and oral surgery and in anesthesiology. Once collected, samples were immediately flash-frozen and stored in liquid nitrogen until analyzed. Tumors were diagnosed as CAA or COSCC using H&E-stained slides reviewed by a board-certified veterinary pathologist based on previously described criteria^41^. To further characterize tumors, *HRAS* p.Q61R genotyping assays were carried out using a PCR and Sanger sequencing approach, as previously described^21^. Only CAA tumors harboring an *HRAS* p.Q61R mutation were included, while only COSCC and healthy gingival tissues harboring *HRAS* p.Q61 wild-type alleles were included.

### Bulk RNA-seq

Samples in DS2 were processed as previously described^16^. For DS1, cryopreserved frozen tissue samples were homogenized in TRIzol (Thermo Fisher) using a bead mill, and the resulting lysate was frozen at -80 °C. After thawing, chloroform was added (20% of total volume) to promote phase separation following the manufacturer’s protocol, After centrifugation, the aqueous (top) phase was transferred to a new tube and an equal volume of 100% ethanol was added. RNA was purified from the sample using a Zymo Quick RNA kit, including treatment with DNAse. RNA concentration was determined with a Nanodrop (Thermo Fisher), and RNA quality was analyzed with a Fragment Analyzer (Agilent). Ribosomal RNA was depleted from 500ng total RNA using the NEBNext HMR rRNA Depletion Kit v2 (New England Biolabs) and RNA-seq libraries were prepared using the NEBNext Ultra II Directional library prep kit (New England Biolands). A minimum of 40M paired-end 2×150 bp reads per sample were generated on a NovaSeq X Plus instrument (Illumina).

### Single nucleus RNA-seq

CAA, COSCC, and healthy gingival samples were homogenized on ice using a Dounce tissue grinder in digestion buffer (10 mM Tris, pH 7.5; 150 mM NaCl; 5 mM MgCl_2_; 1.5 mM CaCl_2_; 0.1% Tween-20; 1 mM DTT; 1 mg/mL Collagenase I; 0.5 mg/mL Collagenase IV; 1× Halt Protease Inhibitor Cocktail; 0.1 U/µL Protector RNase Inhibitor; 0.1 U/µL SUPERase•In RNase Inhibitor). Lysis buffer (10 mM Tris, pH 7.5; 10 mM NaCl; 3 mM MgCl_2_; 0.1% Tween-20; 0.05% NP-40; 1 mM DTT; 0.1 U/µL Protector RNase Inhibitor; 0.1 U/µL SUPERase•In RNase Inhibitor) was added to the homogenate, and the mixture was gently pipetted 10–20 times, and then incubated on ice for 5 min. After lysis, the lysate was centrifuged at 400 × g for 5 min at 4 °C, and nuclei were subjected to multiple rounds of washing with 1× PBS and sequentially filtered through a 35 µm mesh. Nuclei were then fixed according to the Evercode(tm) Nuclei Fixation v3 kit protocol prior to combinatorial barcoding using Evercode WT Mega v3 (Parse Biosciences). Sequencing was performed on the NovaSeq X Plus platform (Illumina).

### Bulk and scRNA-seq data processing

Bulk RNA-seq data was processed as previously described for DS2^16^. For DS1, adapter and low quality bases were trimmed from the 3’ ends and filtered to remove trimmed read pairs with one or more reads < 50 nt in length with Trim Galore v0.6.6/cutadapt v2.8^42^. Trimmed reads were mapped to the CanFam3 reference genome and counts per Ensembl-annotated gene were generated using STAR v2.7.0e^43^. snRNA-seq libraries were processed using standard preprocessing and alignment protocols compatible with the Parse Biosciences data type. Phix reads were first removed and Trim Galore was used to remove low-quality bases and residual sequencing adapters. Prior to alignment, quality metrics were assessed to ensure read integrity. Processed reads were then provided to the Parse Biosciences pipeline (v1.4.1, Parse Biosciences) for barcode demultiplexing, alignment to the canine reference genome (canFam3^44^), and UMI collapsing, resulting in gene-by-cell count matrices. Each library was processed independently, after which the resulting outputs were merged into a single expression matrix. This integration step combined barcoded transcriptomes from all libraries and provided a unified dataset for downstream analyses.

### Quality control for snRNAseq data

All processing was performed using Scanpy^45^. Cells with less than 1,000 total counts or fewer than 500 detected genes were excluded while only genes expressed in over 50 cells were considered. Cells were excluded if they deviated by more than five median absolute deviations for any of the following metrics: i) log total counts, ii) log number of expressed genes, or iii) percentage of counts contributed by the top 20 genes. Furthermore, cells with mitochondrial counts making up over 1% of their total counts were removed. Doublets were identified using scanpy’s integrated Scrublet2, with sim_doublet_ratio set to 2 and a changing expected_doublet_rate. The rate was calculated as the number of cells per well divided by 96×96× # of libraries (8), with a minimum threshold set at 5%. Predicted doublets were discarded and cells with a Scrublet score greater than 0.3 were excluded.

### SnRNAseq postprocessing and clustering

To prepare the expression data for clustering, they were normalized and log transformed with scanpy’s default parameters. Prior to total count regression and gene scaling, the top 2000 highly variable genes were selected using highly_variable_genes and the batch parameter set to the sample level. Principal component analysis (PCA) was performed on the scaled data and followed by sample level batch correction using Harmony^46^. A neighborhood graph was computed using the top 30 batch corrected principal components and n_neighbours set to 100. The uniform manifold approximation and projection (UMAP) embeddings were then calculated and leiden clustering was performed for various resolutions. Patient/sample specific clusters were removed and clusters were annotated as broad cell types using key marker genes from previously published studies. To refine annotations, each broad cell type was individually reclustered to resolve finer subtypes. Throughout the analysis, cells that had hybrid expression patterns of multiple broad cell types were considered as doublets and were discarded.

### Pearson correlation of bulk and pseudobulk dataset DGEA results

To compare the relationship between CAA and COSCC between the snRNA-seq data and bulk RNA-seq datasets, pseudobulk samples were generated using the snRNA-seq data by summing the gene counts per sample. DGEA was assessed on pseudobulked and bulk RNA-seq counts using DESeq2 (via PyDESeq2^47^). For the bulk RNA-seq datasets, genes with counts greater than 5 counts per million (CPM) in at least 5 samples were retained, and DESeq2 was run on the merged datasets using tumor condition and dataset of origin as design factors. For the pseudobulk samples, similarly as before, genes meeting the expression threshold (≥5 CPM in ≥5 samples) were retained, and DESeq2 was run with dispersion estimation, Wald tests, Benjamini–Hochberg false discovery rate (FDR) correction, and log2 fold-change shrinkage to ensure robust and reproducible results.

### Principal component analysis (PCA) for pseudobulk and bulk RNA-seq

Raw bulk or pseudobulk RNA-seq counts were converted to CPM and log-transformed after filtering for genes with >5 CPM in at least five samples. For each analysis, genes were either subset to contain the top 2000 highly variable genes as determined by the Scanpy implementation of HVG selection, or restricted to the CAA-specific KC signature gene set. The genes were then scaled to unit variance and zero mean prior to PCA.

### Pairwise DGEA snRNA-seq data and CAA-specific KC signature generation

Pairwise differential gene expression analyses were performed using the Wilcoxon rank-sum test. Epithelial populations were grouped separately from the TME. The CAA-specific basal KCs were compared one-versus-one against all other epithelial groups and the TME as a whole, excluding CAA-specific junctional KCs. Genes with log_2_FC > 2 and adjusted p val < 0.01 (Benjamini–Hochberg correction) were considered significantly upregulated. Similarly, the CAA-specific junctional KCs were compared one-versus-one against all other epithelial groups and the TME as a whole, excluding CAA-specific basal KCs and the significantly upregulated genes were determined (log_2_FC > 2 and adjusted p val < 0.01). The union of these two DGEAs resulted in the CAA-specific KC signature of 168 genes for downstream analysis.

### MSigDB collection analysis on CAA-specific KC signature

Overrepresentation analysis (ORA) was performed on the CAA-specific KC signature using GSEApy^48^. Gene sets were tested against MSigDB’s C5 (GO gene sets) and C6 (oncogenic signatures) collections and only gene sets with adjusted p < 0.05 (Benjamini–Hochberg correction) were considered enriched.

### RNA isolation and quantitative PCR

Flash frozen biopsy samples from COSCC (n=6), CAA (n=6), and healthy gingiva (n=6) were used for qPCR analysis. RNA was extracted from samples using TRIzol Reagent (ThermoFisher Scientific, Waltham, MA) according to the manufacturer’s instructions, and RNA concentration and purity were determined using a NanoDrop spectrophotometer (ThermoFisher). Samples with 260/280 and 260/230 ratios of approximately 2 were used for the analysis. A total of 500 ng of RNA from each sample was treated with Rnase-free Dnase I (ThermoFisher) and then reverse transcribed into cDNA using the qScript cDNA SuperMix kit (QuantaBio, Beverly, MA), according to the manufacturer’s instructions. Primers for *GAPDH, HPRT1, ERBB4*, and *PEG3* were designed using the NCBI Blast tool or taken from literature, as seen in Supplementary Table 1. Quantitative PCR was run using the QuantStudio 3 real-time PCR system (Applied Biosystems). Total reaction volume was 10 µL, containing 5 µL of PowerTrack SYBR Green Master Mix, 3.6 µL of nuclease-free water, 0.2 µL (200 nM) each of the forward and reverse primers, and 1 µL of cDNA (25 ng). All samples were performed in duplicate along with a no reverse transcriptase control and a no template control. The run included an initial denaturation and enzyme activation step (10 min at 95°C) followed by 40 cycles of denaturation (15 s at 95°C) and annealing (1 min at 60°C). A melt curve was collected from 60 to 95°C to check for non-specific amplification in each well, and Sanger sequencing of the PCR product was conducted at the Cornell Genomics Facility to confirm amplification of the correct product. Expression levels for the target genes were normalized to the mean expression of *GAPDH* and *HPRT1* to obtain Delta Ct values, and relative quantification (RQ) values for CAA were calculated against COSCC.

### IHC staining

Representative formalin-fixed and paraffin-embedded tissue sections obtained from archived normal gingiva (n=4), and dogs with CAA (n=8) or common subtypes of COSCC, including conventional (n=3), papillary (n=3) and basaloid (n=3) were assessed for ERBB4 and PEG3 protein expression by immunohistochemistry (IHC) using standard protocols and an automated IHC stainer (Bond-Max automated IHC staining system; Leica). All samples were submitted as biopsy specimens, obtained between 2023 and 2025. To preserve critical antigenic epitopes, samples that contained bone and had undergone decalcification were not included for IHC. The diagnosis of CAA and COSCC was made based on initial hematoxylin and eosin assessment by board-certified veterinary pathologists. Briefly, 4-µm tissue sections mounted on charged slides were deparaffinized (AR9222, Bond dewax solution; Leica), and after heat epitope retrieval (AR9640, Bond epitope retrieval solution 2; Leica) at pH 9 for 30 minutes, the sections were incubated with PEG3-specific rabbit IgG polyclonal antibody (1:200 dilution; Cat# PA5-99683; ThermoFisher, Waltham, MA, USA) raised against a synthesized peptide corresponding to amino acid residues 1427-1477 of human PEG3, or ERBB4-specific rabbit IgG polyclonal antibody (dilution 1:400; ab137412; Abcam, Cambridge, MA, USA) raised against a proprietary recombinant protein corresponding to amino acid residues 850-1150 of human ERBB4, for 15 minutes followed by polymeric horseradish peroxidase (DS9800; Bond polymer refine detection; Leica) linker antibody conjugate detection system for 10 minutes, and hematoxylin counterstain (DS9390; Leica) for 5 minutes. Positive immunoreactivity was identified as brown staining with diaminobenzidine (DAB). The IHC evaluation was performed independently by G.E.D, while blinded to the results of previous molecular analyses and medical records.

### Kinome regularization (KiR) modeling

KiR models for three CAA cases were generated as previously described^31^. Kinase inhibition profiles of the 32 inhibitors served as explanatory variables, and corresponding microtumor response data were used as response variables in elastic net-regularized multiple linear regression models. Models were built using custom R scripts (https://github.com/FredHutch/KiRNet-Public) with the glmnet package. Leave-one-out cross-validation (LOOCV) was used to select the optimal penalty factor (λ), and models were computed across 11 values of α (ranging from 0 to 1) to balance Least Absolute Shrinkage and Selection Operator (LASSO) and Ridge regularization. Model performance was evaluated using LOOCV error and root-mean-squared error (RMSE) of predictions.

### Microtumor preparation and culture

Microtumors derived from cryopreserved CAA tissues were prepared as previously described^39^. Briefly, tumor slices were generated from freshly thawed specimens using a Leica VT1200S vibratome (Leica Biosystems, Wetzlar, Germany). The tissue was sectioned into slices approximately 400 μm in thickness and 6 mm in diameter. To generate three-dimensional (3D) microtumors (μtumors), the slices were arranged in a monolayer on a sterile plastic disc and sectioned using a McIlwain Tissue Chopper (Ted Pella, Redding, CA, USA). After the initial series of parallel cuts, the disc was rotated 90° and the sectioning process was repeated to create uniformly sized microtumor fragments. The resulting 3D μtumors were cultured in Williams’ Medium E supplemented with 12 mM nicotinamide, 150 nM ascorbic acid, 2.25 mg/mL sodium bicarbonate, 20 mM HEPES, 50 mg/mL glucose, 1 mM sodium pyruvate, 2 mM L-glutamine, 1% (v/v) insulin–transferrin–selenium (ITS), 20 ng/mL epidermal growth factor (EGF), 40 IU/mL penicillin, and 40 μg/mL streptomycin. Microtumors were maintained under standard culture conditions until drug treatment.

### Kinase inhibitor screening

3D μtumors were treated with Realtime-Glo MT cell viability reagent (Promega, Madison, WI, USA) at 1: 1000 dilution in 96-well plates. Following a 24h incubation, luminescence signals were measured using a Biotek Synergy H1 microplate reader. Wells with signal levels at least 10 times higher than the blank were selected for kinase inhibitor screening, with each inhibitor tested in duplicate or triplicate. 3D μtumors were maintained, and real-time viability was measured for 5 days post-treatment. The viability was determined by calculating the ratio of the signal at the end of the study to the signals from the same tissue before treatment, which was then normalized to the value of vehicle control 3D μtumors set to 100%.

## Funding

This work was supported by a Cornell Richard P. Riney Canine Health Center research grant awarded to S.P and W.P.K., a philanthropic gift from the Joseph and Bessie Feinberg Foundation awarded to P.S., a philanthropic gift from the Morgan Cueman Cancer Research Fund awarded to P.S. and S.P.,and a grant from the National Science Foundation (NSF Award No. 2047289) awarded to T.S.G.

## Data availability

The authors declare that all RNA-seq data generated in this study have been deposited in NCBI’s Gene Expression Omnibus (GEO) with GEO series accession numbers GSE317244 (bulk RNA-seq) and GSE317641 (snRNA-seq). RT-qPCR results are available as supplementary data.

## Code availability

All code needed to reproduce the bulk and snRNA-seq analysis presented in this study have been deposited on Github: https://github.com/AndreasStephanou/CAA_Paper

## Competing interest statement

The authors declare no conflict of interest.

## Supplemental Figures

**Supplemental Figure 1.**
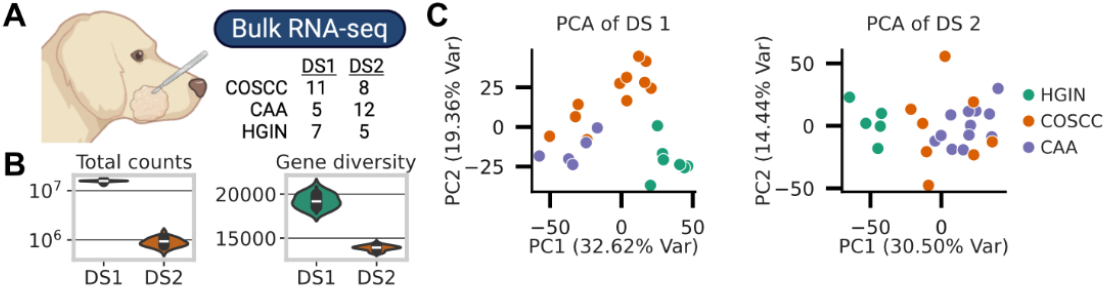
**A)** Experimental workflow of CAA, COSCC, and HGIN bulk RNA-seq. Created with BioRender.com. **B)** Violin plots showing the i) total counts (reads), and ii) number of non-zero expression genes across samples per dataset. **C)** PCA plots of bulk RNA-seq samples using the top 2000 most highly variable genes per dataset.

**Supplemental Table 1.**
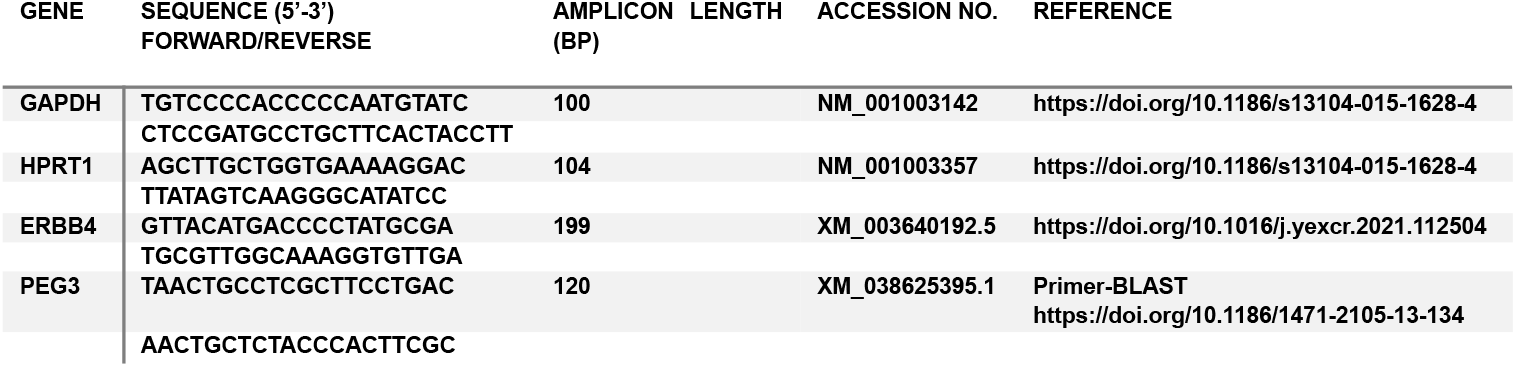
Primer sequences for genes used for RT-qPCR.

## Notes

### Competing Interest Statement

The authors have declared no competing interest.

